# Biochemical noise enables a single optogenetic input to control identical cells to track asymmetric and asynchronous reference signals

**DOI:** 10.1101/2022.07.05.498842

**Authors:** Michael P May, Brian Munsky

## Abstract

Optogenetics is a powerful technology to control synthetic gene circuits using external and computer-programmable light inputs. Like all biological processes, these systems are subject to intrinsic noise that arises from the stochastic process of gene regulation at the single-cell level. Many engineers have sought to mitigate this noise by developing more complex embedded bio-circuits, but recent work has shown that noise-exploiting stochastic controllers could enable new control strategies that take advantage of noise, rather than working against it. These noise-exploiting controllers were initially proposed to solve a single-input-multi-output stationary control problem, where symmetry was broken in a means reminiscent to the concept of Maxwell’s Demon. In this paper, we extend those results and show through computation that transient, asymmetric, and asynchronous stochastic control of the single-input-multi-output (SIMO) control problem is posible to achieve by cycling through different controllers in time. We show that such a method is able control two cells to two different periodic fates with different frequencies and different phases despite the use of only one control input.

## I. Introduction

The combination of synthetic biology, fluorescence labeling, optical microscopy, and computer controlled optogenetics has created the possibility for cyber-organic control mechanisms that watch cellular behaviors in real time and then actuate internal cellular processes through application of external light inputs [1], [2]. Such circuits could enable many potential applications that require precise spatial and temporal control of individual cells, but unique challenges arise from the unique spatiotemporal scales of the biochemical reactions that define cellular process. Most importantly, biological behaviors are governed by fundamentally stochastic processes due to small numbers of DNA, RNA, and proteins that interact in discrete and randomly timed events – these events can create heterogenous responses within a population of isogenic cells [3]. Because the regulation of genes in synthetic biocircuits also exhibit such behavior, stochastic gene regulation has been treated as a fundamental limit to the controllability of biological systems. Many have tried to build biological systems that reduce noise [4], but recent work has shown that noise can create new control opportunities [5], [6].

Although many models for optogenetically controlled synthetic cell dynamics have utilized ordinary differential equations, linear noise approximations with additive noise, or Monte Carlo methods such as Gillespie’s stochastic simulation algorithm [7], improved computational tools are still needed to improve understanding for these dynamics and to assist efforts to mitigate or exploit randomness in stochastic gene regulation. In this paper, we use direct analyses of the full probability distributions of system responses under transient inputs (i.e., solutions to the chemical master equation) to find deeper understanding for the stochastic cellular process. We show that this analysis can support the design of controllers that can improve performance of optogentic systems including for SIMO control. We show how the combination of optogenetics, fluorescent microscopy, and noise-exploiting feedback control could yield new control techniques which could able the control of two cells to different periodic fates.

The integration of stochasticity and feedback to stretch the limits of physical possibilities extends back to 1867, when James Maxwell proposed a theoretical demon who could violate the 2nd law of thermodynamics to create a temperature gradient without consuming energy. In essence, the demon uses the random transient state of the system, combined with a controllable trap door to direct fast (hot) molecules to one side and slow (cold) molecules to the other, despite the fact that all molecules sample form the same velocity distribution. In previous work, we proposed a similar optogenetic Maxwell’s demon (OMD) that exploits the transient nature of biological noise in order to direct two identical cells to two different aribitrarily chosen phenotypes. The unique challenge of this control is that the demon cannot just “open the door for a single cell”, but rather if the demon changes the light to control one cell, that same light signal will affect and potentially destabilize the other Fig1(A). The demon must play a difficult balancing game to drive each cell to a different phenotype simultaneously without destabilizing the other.

**Fig. 1.**
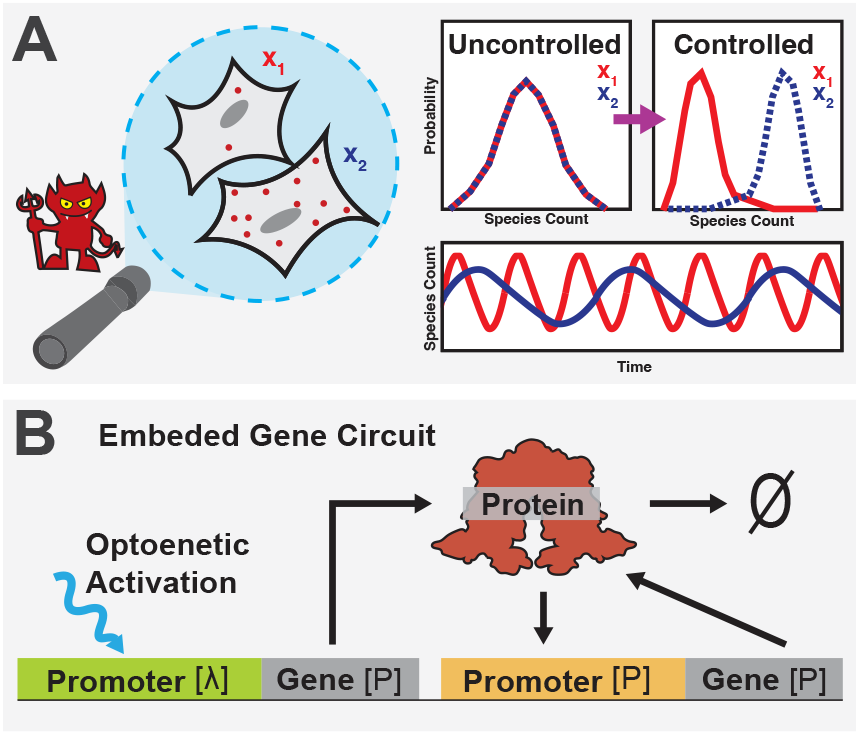
(A). Shows an OMD which seeks to control a stochastic two cell system to two different reference points despite the fact that they both share the same embedded gene circuit and global light input. (B). Schematic of a non-linear and optognetically controllable embedded gene circuit for the production of an arbitrary protein.

Although this previous work performed well at breaking the symmetry between the two cells, our previous work only considered the stationary behavior about a single static reference point. When the control of a system to a periodic process is desired [8], such a control method may not be effective. In this paper, we extend the analysis of previous work to enable the stochastic control toward dynamically changing reference points Fig1(A).

In the next section, we introduce a nonlinear stochastic model for two-cell SIMO control, and we demonstrate how to compute the stationary and transient evolution of probability mass under different control laws. Next, we introduce a metric to evaluate the probabilistic performance error of this controller and devise a gradient based search to optimize the controller to minimize this error. In Section III, we show our results that for the tracking of different dynamic reference signal where two cells are forces to follow different asymmetric and asynchronous trajectories. Finally, in the Discussion Section, we summarize our results and discuss current challenges and future efforts to control multiple different cells using a limited number of control inputs.

## II. Methods

### A. Nonlinear Model for Noise-Exploiting Feedback Control

We previously showed that the combination of stochastic single-cell variation, nonlinear gene regulation, and global feedback has the potential to break the symmetry between identical cells and enable the individualized control of multiple cells using a single control input [5]. One possibility for such a system is illustrated in Fig1(B), which depicts a primary actuator promoter that allows for external feedback control and a secondary self-actuated promoter that provides a nonlinear stabilization once the system is activated. Although the primary promoter in the model to be described in this manuscript is based on the optogenetic T7 dimerization system from Baumschlager et al, [1], whose model parameters we fit in previous work [5], other actuator promoters could utilize alternative electronic, optical, or chemical signals to induce gene expression.

To model this system, we assume that the total protein production rate, *H*(*x*), due to the two promoters is defined by:

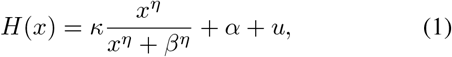

where *x* is the number of protein molecules; *κ, η* and *β* are the max-value, Hill coefficient, and half-max activation concentration, respectively, for the self-actuated promoter; *u* is the strength of the optogenetically controlled promoter; and *α* is the total basal (or leaky) expression level from the two promoters. The protein decay rate is assumed to be a first order reaction with rate:

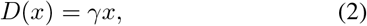

where *γ* represents the degradation rate. Specific values for all parameters are provided in Table I.

**TABLE I.** Parameters of the Model

| Parameter Name | Value | Units |
| --- | --- | --- |
| $\kappa$ | 0.406 | molecules minute<sup>-1</sup> |
| $\beta$ | 20.0 | molecules |
| $\eta$ | 8.00 | |
| $k_0$ | 0.0001 | molecules minute<sup>-1</sup> |
| $\gamma$ | 0.0203 | minute<sup>-1</sup> |

For the discrete stochastic description of such a system consisting of two cells, let *x*_1_ denote the integer protein population in the first cell, and let *x*_2_ denote the protein population in the second cell. In this case, there are four possible production events (one for each promoter in each cell) and two degradation events (one for each cell). The stoichiometry for these six possible events can be written in matrix form as:

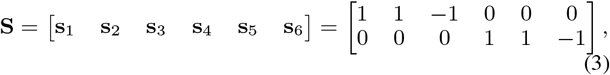

and the corresponding propensity functions (i.e., the stochastic reaction rates) for these reactions are given by:

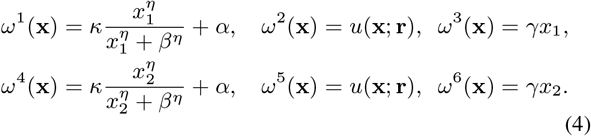

The propensity functions for the optogenetic promoter, *ω*^2^ and *ω*^5^, are identical due to the fact that a single light-modulated control signal is applied to both cells at the same time. The control signal, *u*(**x**; **r**), can be any positive valued function of the current state 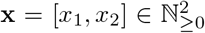 and some reference signal 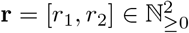. In other words, *u* maps 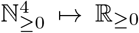. We stress that the control signal *u*(**x**; **r**) is equal for both cells due to the fact that the global light input that shines on both cells at once and gives rise to the SIMO control system as shown in Fig1(A). The goal of the stationary OMD from [5] was to find an optimal *u*(**x**; **r**) to direct the discrete stochastic process 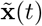 to one specific reference point **r**. In this manuscript, we explore the possibility to allow **r** to vary as a function of time.

### B. Chemical Master Equation description for the joint dynamics of two cells under a single control input

The state space for the two-cell process introduced above is the set of all non-negative integer vectors 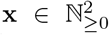. Using the stoichiometry vectors {**s**_**1**_, …, **s**_**6**_} and propensity functions {*ω*^1^(**x**) …, *ω*^6^(**x**) } defined above, we can state the chemical master equation (CME [9]) to quantify the time rate of change for the probability of each state **x** as:

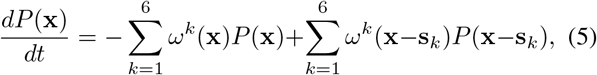

where the first sum describes flow of probability away from state **x** and the second sum describes flow of probability into state **x** from its neighbors. We can enumerate the countable state space **x** ∈ ℕ_*≥*0_ according to: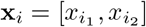, where the *i* = *i*(*i*_1_, *i*_2_) is an arbitrary enumeration scheme to be defined below. Under any such enumeration scheme, we can define the probability mass vector: **P** = [*P* (**x**_1_(*t*), *P* (**x**_2_(*t*), …]^*T*^, and write the CME in linear matrix form as:

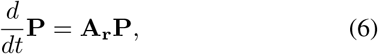

where the infinitesimal generator, **A**_**r**_, for a specific reference point **r** is given by:

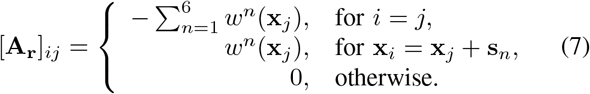

Although there is an infinite number of possible states, **x** ∈ ℝ_*≥*0_, one can approximate the CME solution using a truncation such as the Finite State Projection approach [10] for finite times or by adding a reflecting boundary condition for use with steady state analyses. Specifically, for the purposes of this manuscript, we truncate the state space such that *x*_1_ *< N* and *x*_2_ *< N*, which allows us to use the relatively simple enumeration scheme:

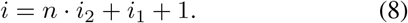

For all analysis presented here, *n* = 50, such that the number of possible states (and therefore the length of the vector **P**) is (*n* + 1)^2^ = 2500.

To approximate steady state of the CME, we assume that the propensity functions *ω*^1^ and *ω*^2^ are both zero when *x*_1_ = *n* − 1 and that *ω*^4^ and *ω*^5^ are both zero when *x*_2_ = *n* − 1, such that the number of proteins is not allowed to exceed *n* molecules in either cell. Under this approximation, the restricted steady state probability distribution can be estimated first by computing the null space of the infinitesimal generator,

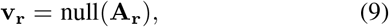

and then normalizing this result to have a sum of unity:

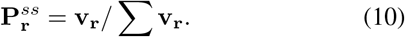

In the above formulation, 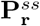 should be understood as the joint steady state probability distribution over the enumerated state space 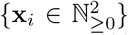 under a fixed control law *u*(**x**; **r**). We will later use **P**_**r**(*t*)_(*t*) to denote the solution of the master equation under a potentially time-varying set of control laws, *u*(**x**; **r**(**t**)). In either case, the corresponding marginal distributions *P* (*x*_1_) or *P* (*x*_2_) can be calculated by using a linear operations:

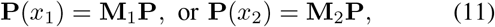

where **P** can be replaced with 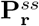 or **P**_**r**(*t*)_(*t*) and for the appropriate linear marginalization matrix **M**_1_ or **M**_2_.

### C. Metric to Quantify Control Performance Error

To rank outcomes of different control laws, we use the expected the mean **squared** error between the cell state and the reference signal,

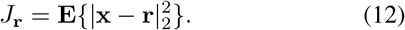

This score is calculated by summing over all possible states:

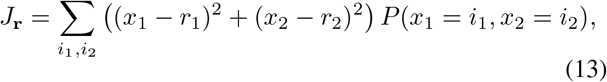

which can be written more concisely in terms of the enumerated state space:

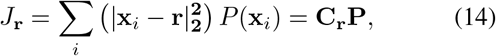

where the *i*^th^ entry of the reference-dependent output vector is given by 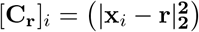.

### D. Optimizing the Control Law to Minimize Error

Using the definitions above, we can now optimize to find a control law *u*(**x**; **r**) that minimizes the mean squared error for any specific reference point **r** (i.e., to minimize *J*_**r**_). We assume that all control signals must be constrained such that:

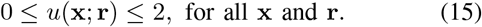

For any given fixed **r**, this optimization problem requires finding a vector of control values **u**_**r**_ = [*u*(**x**_1_; **r**), *u*(**x**_2_; **r**), …] with one entry for every **x**_*i*_ (i.e., for 2,500 different states in our case). The optimization is conducted using a gradient search, where the effect of the control on the error is:

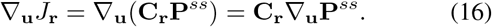

From Eq. 6 at steady state, we can write:

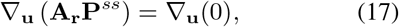

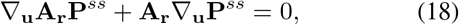

from which we calculate ∇_**u**_**P**^*ss*^ using the generalized minimal residual method [11].

To estimate an optimized solution, we begin with some initial control law **u**_0_ and follow the gradient descent procedure according to:

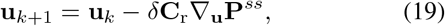

where *δ* is a small learning rate. The search defined by Eq. 19 is looped over many iterations *k* = 1, 2, …, where on each iteration **A**_**r**_ and ∇_**u**_**A**_**r**_ are updated according to Eq. 7; then **P**^*ss*^ is found using Eqs. 9 and 10; and ∇_**u**_**P**^*ss*^ is determined from Eq. 18. Finally, after the optimization converges, the control inputs are bounded such that any **u** greater than **u**_*max*_ is set to **u**_*max*_.

### E. Control to follow a path of changing reference points

For a single fixed reference point **r**, the optimization above performs well to find a locally optimal controller that minimizes a static *J*_**r**_ for that specific reference state. For systems where a changing reference point may be desired, a single reference point solution is no longer sufficient. To extend these results for a dynamic reference signal, we compute the locally optimal steady state controllers at several discrete points along the desired pathway.

For our examples below, we restrict the reference signals to periodic functions with a period *T*, although non-periodic functions could be considered using the same approach. The piecewise constant definition of the reference signal is given as

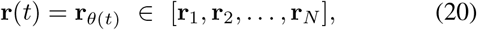

where the index, *θ*(*t*) is defined as:

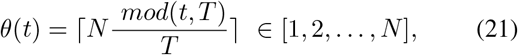

and the notation rounds ⌈·⌉ its argument up to the nearest integer. The corresponding piecewise constant controller **u**(*t*) is then defined by selecting the appropriate reference value at the appropriate time:

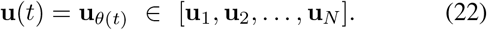

Finally, to solve for resulting time-inhomogeneous CME, we simply integrate:

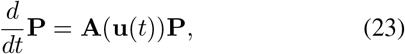

but where the infinitesimal generator matrix (Eq. 7) is now a piecewise constant function in time.

As a result of the dynamic changes to the reference point, the resulting probability distribution **P**(*t*) = **P**(**x**, *t*) changes continually in time and no longer reaches a fixed steady state value. To visualize the error of the control performance, we define the random error 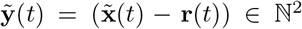 and translate the computed probability distribution to calculate the joint distribution of the error at any specific instance in time as:

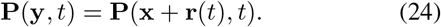

The best possible controller is one where most of the probability mass concentrates at **y** = [0, 0], and where very little probability mass accumulates at large magnitudes of **y**. In order to visualize the average error (direction and magnitude) over a periodic trajectory, we integrate to average over time:

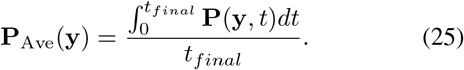

## III. Results

Optimized control inputs were found on a square grid of static reference points which minimized the score 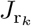 at each point in a coarse grid. Each element of the grid *G* ∈ [*r*_1_, *r*_2_] is created by stepping *r*_1_ and *r*_2_ from zero to forty in steps of five for a total of 9^2^ = 81 grid points. Optimal scores 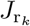 were found for each static point in the grid and are plotted on the domain of grid points as a heat map Fig 2(A). The heat map shows that some static reference points in the domain are more easily controlled than others, with lower scores (lighter colors) indicating more controllable static reference values and higher scores (darker colors) indicating worse control. Higher scores are seen in the top-left, bottom-right, and the middle region of the heat map Fig 2(A). The edge regions likely have poor control due to the geometry of the system, while the middle point **x** = [20, 20] has poor control due to instability of the rate equations for that particular value of **x**. The dashed black line in Fig 2(A) shows a line of symmetry in the score values. Transposing the heat map along this line yields a perfectly symmetric result because transposing the control matrix is the same as swapping *u*(**x**, [*r*_1_, *r*_2_])*x* → *u*(**x**, [*r*_2_, *r*_1_]).

**Fig. 2.**
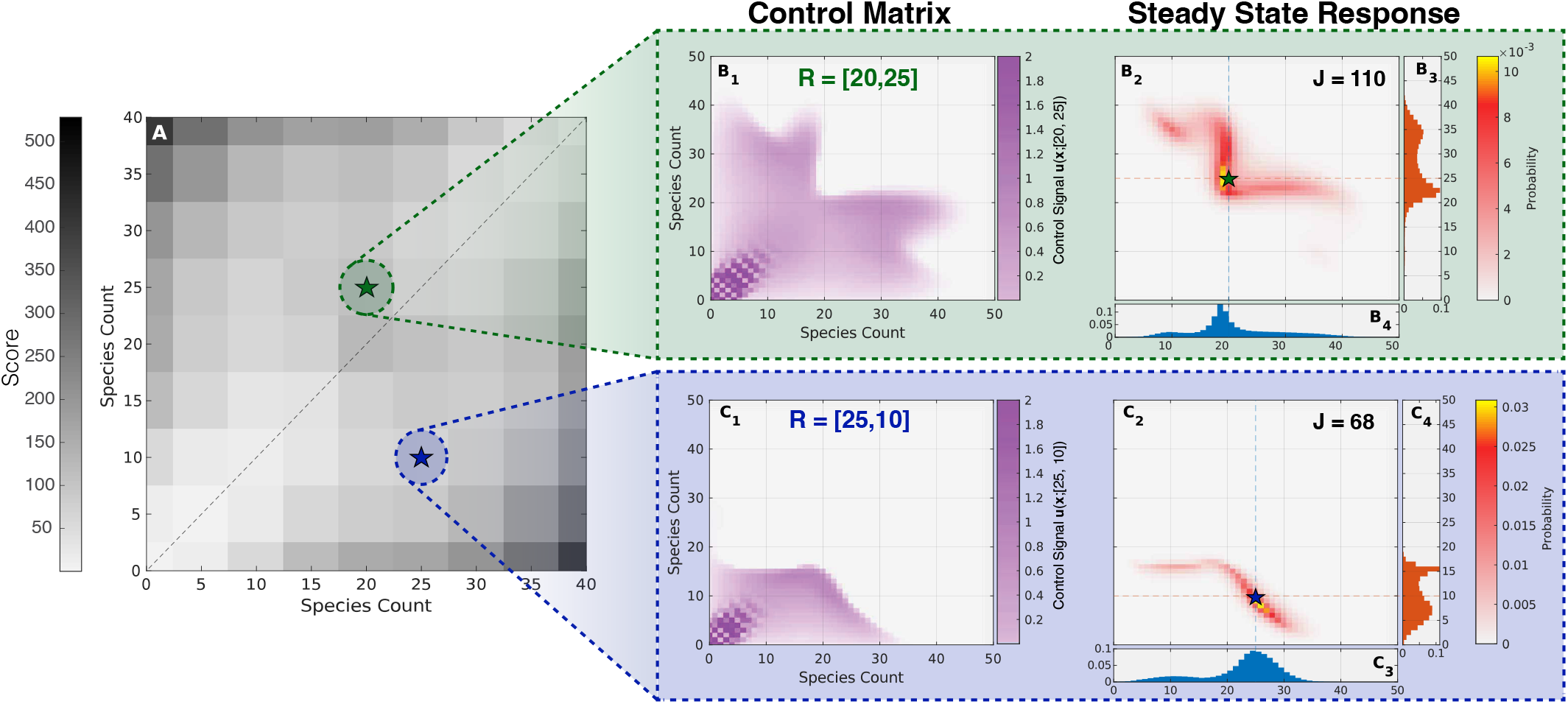
Control laws and performance for fixed reference points. (A) Optimized scores *J*_**r**_ for on a 2D domain with a grid of reference points. Two specific reference points are selected (**r** = [20, 25], green highlighted star) and (**r** = [25, 10], blue highlighted star). (B1) Optimized control law *u*(**x**, [20, 25]) for the desired reference point [20, 25]. (B2) Steady state joint probability distribution of the system when using the control input law depicted in B1. (B3 and B4) Marginal distribution **P**(*x*_1_) and **P**(*x*_2_), calculated using the joint distribution in B2. (C1-C4) Same as for B1-B4, but for reference point of [25, 10].

Figures 2(A, B1, and C1) shows the optimized control laws (*u*(**x, r**)) for two examples of static reference points: **r** = [20, 25] (green highlighted star) and **r** = [25, 10](blue highlighted star). Figures 2(B2 and C2) shows the stationary joint probability distributions under those fixed-reference controllers. These figures show that controllers optimized to different fixed reference points required very different control laws and result in qualitatively different joint probability distributions. The corresponding marginal distributions depicted in Fig 2(B3,B4,C3,C4) show how symmetry between the two cells has been broken despite using only a single global light input signal and for when both cells share identical gene circuits. Joint and marginal distributions of the responses reveal that tighter control of the probability distribution leads to lower score for the more asymmetric reference point (*J*_[25,10]_ = 68 in Fig 2.C2) compared to the more symmetric reference point (*J*_[20,25]_ = 110 in Fig 2.B2).

Using different combinations of the fixed-reference control laws, we specified three controllers to track periodic reference signals:

1. a synchronized trajectory given by

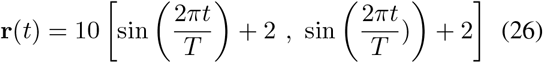

where the desired reference points are the same;
2. a phase lagged trajectory given by

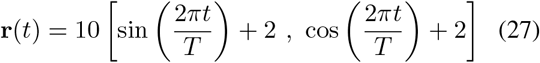

where the desired two reference points contains a *π/*2 phase shift between each other; and
3. a phase lagged trajectory with a two different frequencies

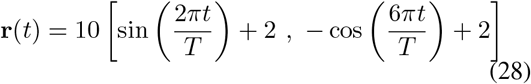

where the second cell is expected to fluctuate with three times the frequency as the first and with a phase shift.

For each of these 2D reference trajectories, *N* = 32 points along the pathway are selected and 32 different controllers are optimized for each point on the path. The period is set to *T* = 5000 minutes. For each reference trajectory, the system is analyzed for a total of five periods, where only the responses over last four periods are shown in each of the plots.

The synchronized reference signal (Fig. 3, Column A) consists of a symmetric reference signal, *r*_1_(*t*) = *r*_2_(*t*). Fig 3(A4) shows **P**(*x*_1_, *t*) (red cloud) and **P**(*x*_2_, *t*) (blue cloud) and the median of the probability distributions over time of *x*_1_ and *x*_2_ (red and blue lines) using the pathway control method. The appearance of a purple probability cloud in Fig 3(A4) is due to the overlap of **P**(*x*_1_, *t*) and **P**(*x*_2_, *t*). The similarity between the median in Fig 3(A4) and the reference points Fig 3(A3) suggest that the system was able to be controlled to the dynamic reference signal, although with some error due to the stochastic nature of the process. Fig 3(A5) shows the root-mean-squared-error (RMSE) given by the square root of *J*_**r**(*t*)_(*t*) versus 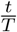. The spikes in the RMSE occur at the points in time when the reference signal crosses through an unstable point of the system’s macroscopic dynamics. This point, where *r*_1_ = *r*_2_ = *β* = 20 (i.e., at the half maximal state of the self-actuating promoters Hill function in Eq. 1) was also seen to be poorly controllable for fixed reference signals, and shows up as a dark patch at that location in Fig 2(A).

**Fig. 3.**
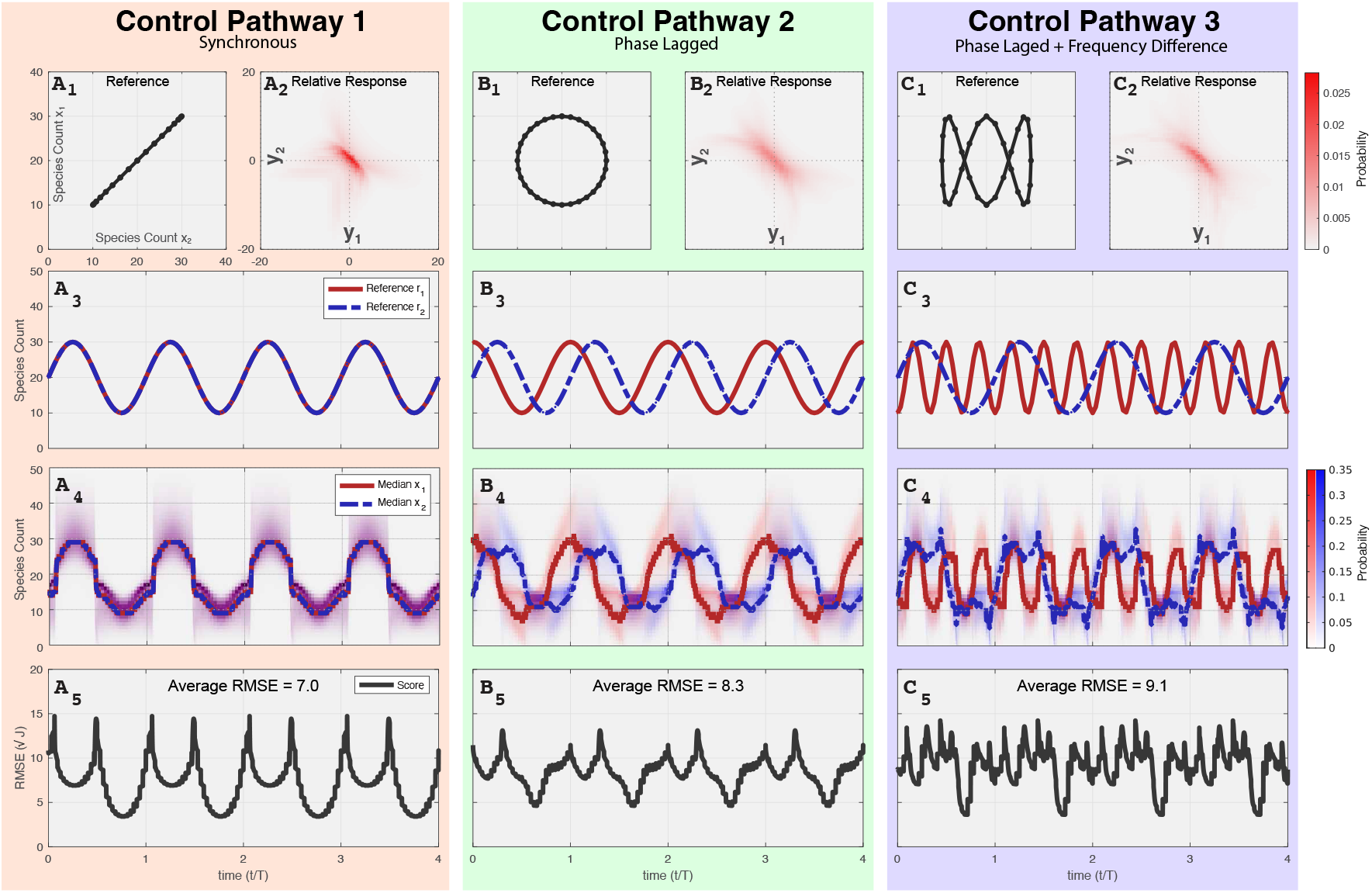
(Control performance for transient reference signals. Three signals are considered: (Column A) synchronous (*r*_1_ = *r*_2_ = 10 sin (2*πt/T*)+20), (Column B) asynchronous with *π/*2 phase lag (*r*_1_ = 10 sin (2*πt/T*) + 20, and *r*_2_ = 10 cos (2*πt/T*) + 20), and (Column C) asynchronous and aperiodic (*r*_1_ = 10 sin (2*πt/T*) + 20, and *r*_2_ = − 10 cos (6*πt/T*) + 20). These reference signals are shown in (A1,B1,C1) 2D phase space (*r*_1_ vs. *r*_2_) and (A3,B3,C3) each versus time. (A2,B2,C2) Time-averaged joint distribution of error (Eq. 25) shows that the response remains well centered about the reference. (A4,B4,C4) Median of *x*_1_ (red line) and *x*_2_ (blue line) show that the controlled periodic behavior matches well to the reference signal. FSP analyses show that probability distributions *P* (*x*_1_, *t*) (red cloud) and *P* (*x*_2_, *t*) (blue cloud) are densely-distributed about the reference trajectory. (A5,B5,C5) Plots of root-mean-square-error versus time; the time-averaged RMSE is shown at the top of each plot.

This process was repeated for the circular (phase-lagged) reference trajectory where *r*_1_(*t*) = 10(sin(2*πt/T*))+20, and *r*_2_(*t*) = 10(cos(2*πt/T*)) + 20. For the **P**(*x*_1_, *t*) (red cloud), **P**(*x*_2_, *t*) (blue cloud) and medians of each (red line, blue line), we find that the solutions generally match the shape and phase lag of the specified reference signals. The data taken together suggest that it is indeed possible to commit two cells to different periodic fates using noise-exploiting feedback control and a single control input. However, we note that the error is not a simple shift from one cell to the other. Rather, the error in cell 1 using clockwise motion matches that of cell 2 in a counter lock wise motion. This effect is clear also in the RMSE error (Fig 3B4) which now spikes four times per period; twice when *r*_1_ = *β* and twice when *r*_2_ = *β*.

Finally, the analysis is repeated on a system where the two dynamic reference points have different phases and frequencies given by *r*_1_(*t*) = 10(sin(2*πt/T*)) + 20, and *r*_2_(*t*) = − 10(cos(6*πt/T*)) + 20 (Figs 3C1,C3). For this system, **P**(*x*_1_, *t*), **P**(*x*_2_, *t*) and the medians are calculated in similar fashion. Looking at Fig 3(C4), we can see that the frequency of the median of *x*_1_ is three times the frequency for *x*_2_, and they generally match the phase and frequency of the desired reference points. Therefore, we can confirm that two cells can generally be controlled to two different frequencies and different phases using the control method, despite the systems being identical and using only a single global light input. The RMSE error (Fig 3C4) now now spikes eight times per period; twice when *r*_1_ = *β* and six times when *r*_2_ = *β*.

Plots of **P**_*Ave*_(**y**) in each of the three controllers (Figs 3A2,B2,C2) show that the synchronous pathway seems to have the most probability mass centered near the origin. This controllability in the synchronous pathway can also be confirmed with the RMSE plotted versus time in (Figs 3A5,B5,C5). The plots of **P**_*Ave*_(**y**) for the asynchronous trajectories (Figs 3B2,C2) show more probability mass at larger magnitudes of error, and these results can be quantified by the higher average RMSE of 8.3 and 9.1 respectively.

## IV. Discussion

Our results show that a noise-exploiting controller is theoretically capable of controlling a stochastic two cell system to different asynchronous and aperiodic trajectories. To design these controllers, we derived a master equation analyses where the control single input was a function of the current random state of the observed cells 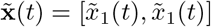 and a temporally changing reference signal **r**(*t*) = [*r*_1_(*t*), *r*_1_(*t*)]. We note that ordinary differential equations could not have been used to model this system, as stochasticity is critical to break the system symmetry and enable control.

The controllers discussed above exclusively were optimized assuming steady state reference points and then modified to reflect the changing reference signal. Unfortunately, it takes for the CME to reach steady state upon changes in the control law, so such controllers may not be optimal for trajectories with faster periods. It would be interesting implement controllers to exploit the transient nature of the CME dynamics. For example, one could envision adaptive controllers based on counterdiabatic driving [12] to temporarily sacrifice steady state responses to drive faster transient changes.

